## Supplementary material for "Sequential Membrane Remodeling by Cholesterol Distinctly Modulates HCN Channels in Naïve and Neuropathic DRG Neurons": This file includes Supplementary Figs 1 - 7 and Table S1

This file includes Supplementary Figs 1 - 7 and legends

**Figure S1**

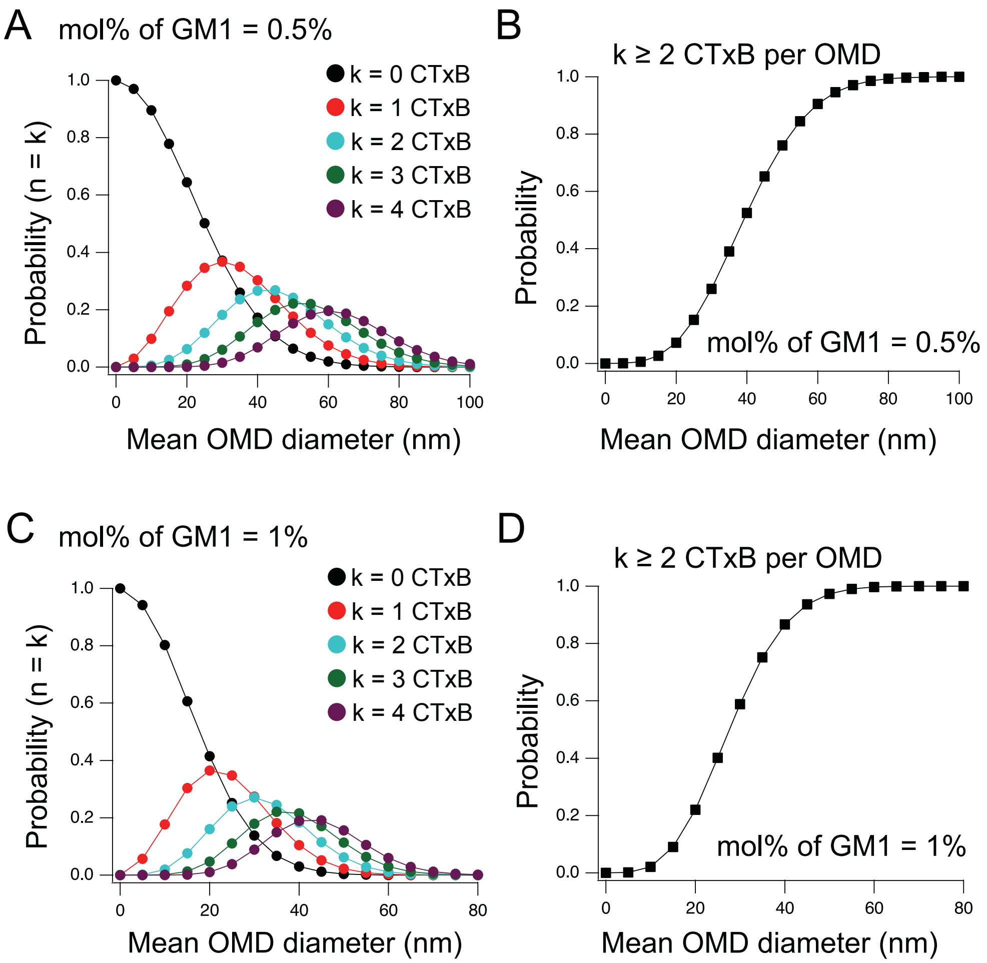


**Fig. S1. Poisson statistics–based simulations illustrating the increased probability of CTxB binding as OMD size increases.** Relative to simulations using 2 mol% GM1 (Fig. 2), simulations shown here assume lower GM1 densities within OMDs: 0.5 mol% (A, B) and 1 mol% (C, D). Within a given cell type, GM1 density within OMDs is assumed to remain constant.

**Figure S2**

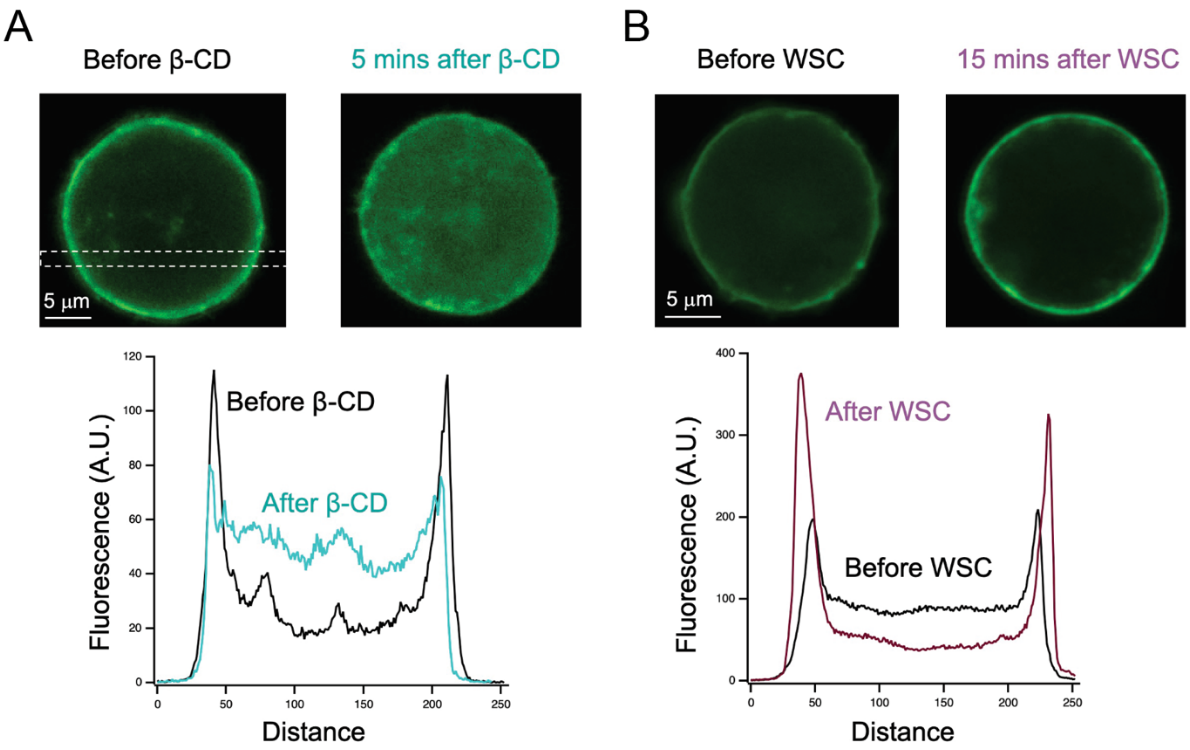


**Fig. S2. Confocal imaging showing the effects of cholesterol extraction or supplementation on membrane localization of GRAM-W-eGFP.**

(**A**) tsA cells overexpressing GRAM-W-eGFP, before and after 5 mins treatment of 2.5 mM β-CD. Line scans of an area such as that indicated by white dashed lines are plotted to highlight the increase in cytosolic GFP fluorescence after β-CD. (**B**) tsA cells overexpressing GRAM-W-eGFP, before and after 15 mins treatment of 0.5 mg/mL WSC, highlighting the increase in membrane-localized GFP fluorescence after cholesterol supplementation.

Figure S3

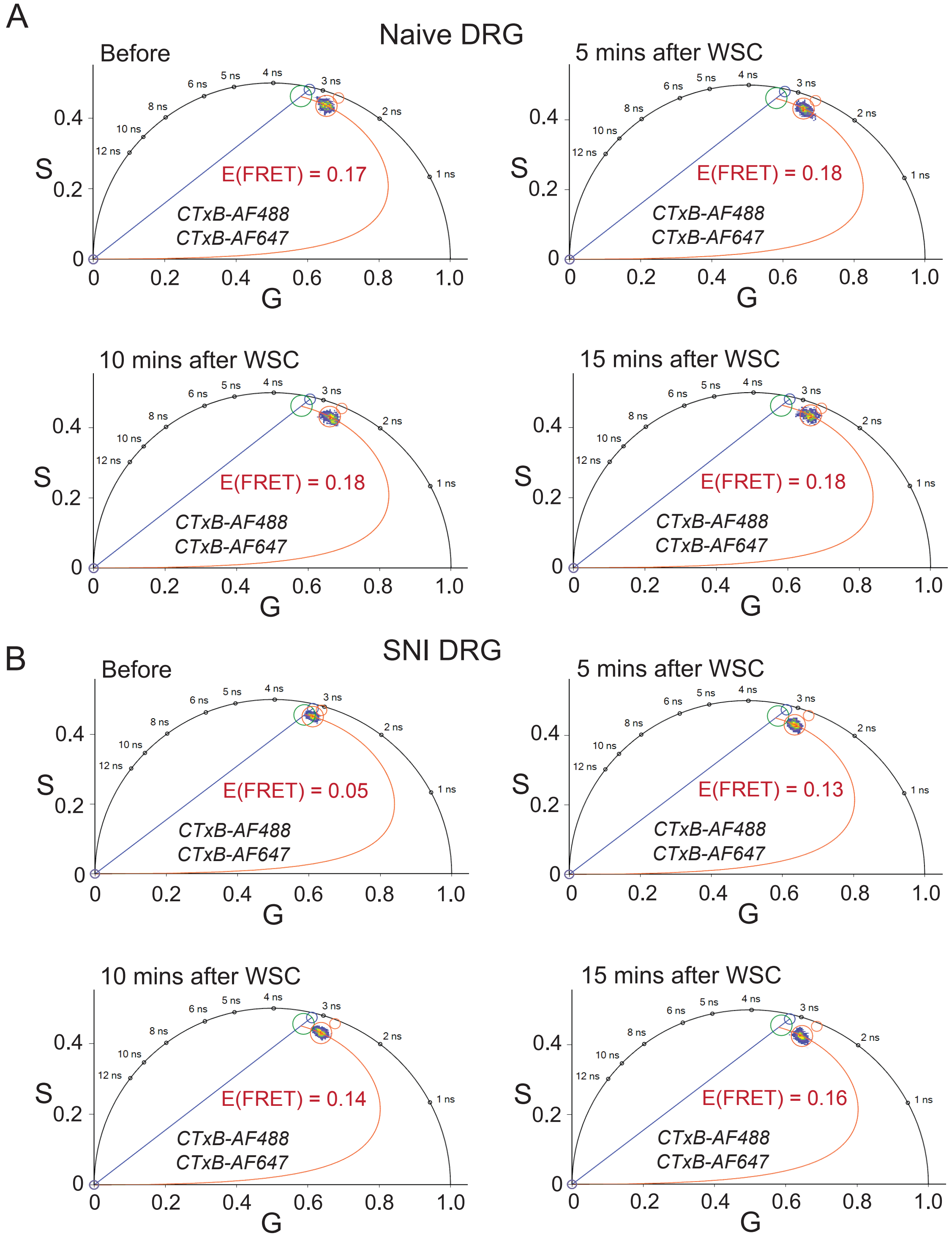


**Fig. S3. CTxB-based FLIM-FRET responses to cholesterol enrichment in naïve and SNI nociceptor DRG neurons.**

**A–B.** Representative phasor plots showing membrane-localized fluorescence in naïve (A) and SNI (B) nociceptor DRG neurons labeled with CTxB AF-488 and CTxB AF-647, recorded before and at 5, 10, and 15 minutes after water-soluble cholesterol (WSC) treatment.

**Figure S4**

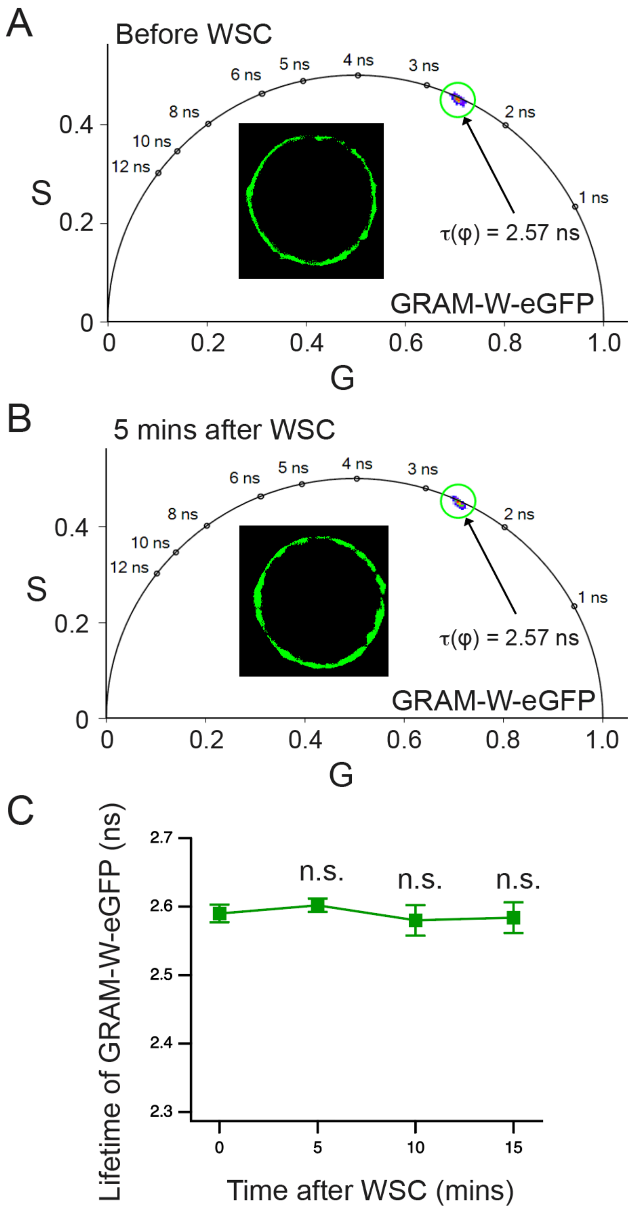


**Fig. S4. WSC treatment does not change the fluorescence lifetime of eGFP-GRAM-W.**
**A-B**. Phasor analysis of eGFP-GRAM-W at the plasma membrane of tsA cells before and 5 min after WSC treatment. Plots depict fluorescence distribution and the corresponding apparent phase lifetime ($\tau\varphi$). **C**. Time course of the mean phase lifetime ($\tau\varphi$) shows no change over 15 min of WSC treatment (mean ± s.e.m.; n = 5 cells; n.s., not significant using paired t-test). During this period, eGFP-GRAM-W exhibited a decrease in fluorescence anisotropy and an increase in intensity, consistent with sensor clustering and increased local density. In contrast, restriction of GFP mobility due to aggregation would be expected to increase anisotropy. Moreover, the stability of the fluorescence lifetime under these conditions rules out a change in rotational mobility as the cause of anisotropy decrease. Instead, these data are indicative of depolarization via an increase in homo-FRET efficiency.

**Figure S5**

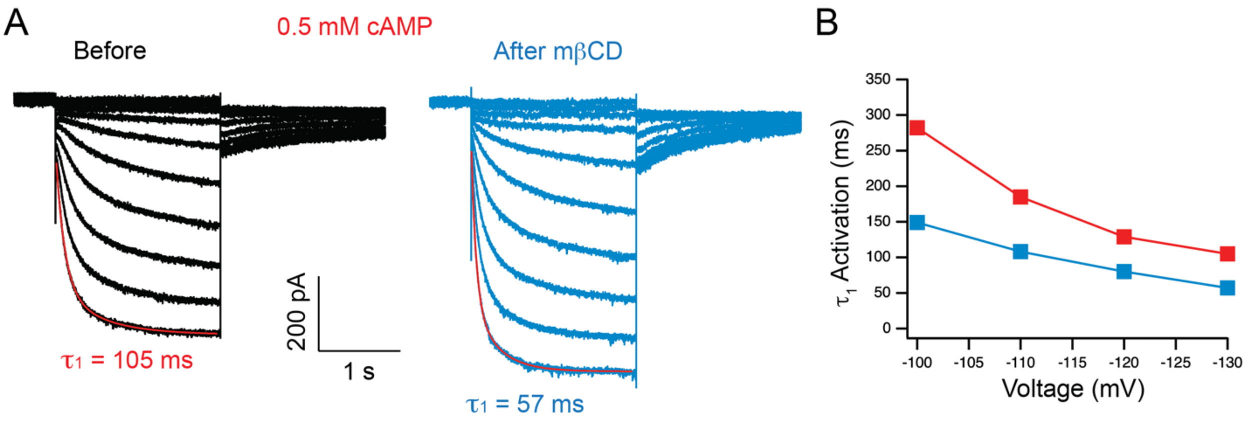


**Fig. S5. Increase in current amplitude and acceleration of the HCN channel activation by cholesterol depletion in the presence of a saturating concentration of cAMP.**

**A.** Representative whole-cell HCN current traces recoded from naïve nociceptor DRG neurons in the presence of the added 0.5 mM cAMP in the patch-clamp pipette solution. Current amplitude increased and the kinetic of channel activation was faster. **B**. Measured primary time constant of channel activation at different hyperpolarizing voltages from the same patch in the panel A.

**Figure S6**

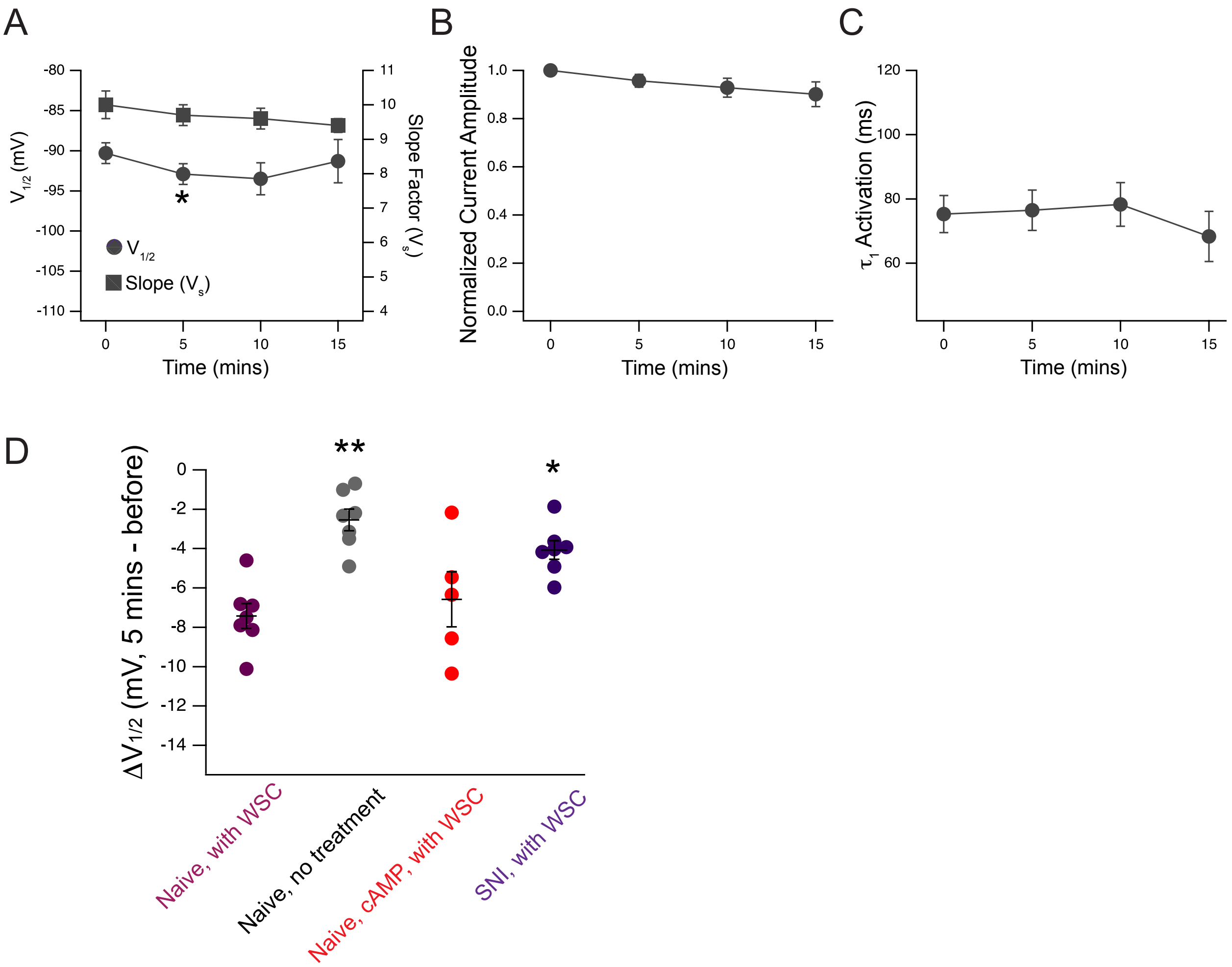


**Fig. S6. Control experiments of the modulation of HCN channel gating for naïve nociceptor DRG neurons without WSC treatments.**

(**A-C**) Summary time course of the change in the $V_{1/2}$ and slope factor ($V_s$) (A), the current amplitude (B), and $\tau_1$ of channel activation (C) post 0.5 mg/mL WSC application for naïve DRG neurons. Data shown are mean ± s.e.m., n = 7 patches. (D) Summary of the change in $V_{1/2}$ ($\Delta V_{1/2}$) 5 mins after with or without WSC treatment for naïve and SNI DRG neurons. Added 0.5 mM cAMP in the pipette solution was also included as a comparison, showing no difference from the condition without the added cAMP, suggesting the effect was mediated by WSC. Data shown are mean ± s.e.m., n = 5-7 patches, one-way ANOVA, $*p < 0.05$, $**p < 0.01$.

**Figure S7**

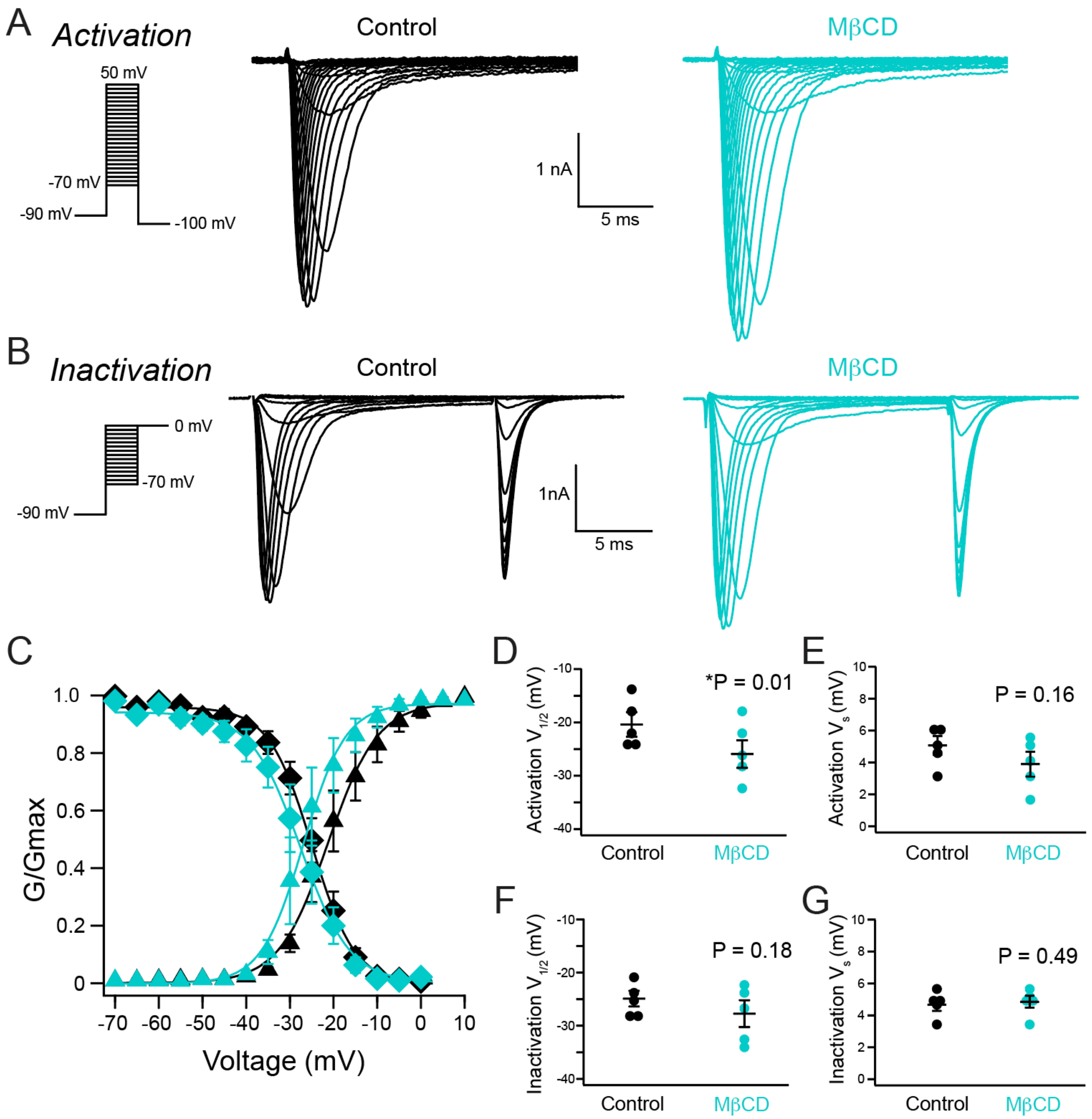


**Fig. S7. Effects of cholesterol extraction on the gating of endogenous sodium currents of naïve nociceptor DRG neurons.**

(**A-B**) Representative currents caused by sodium channel activation (A) and inactivation (B) in naïve nociceptor DRG neurons, before and after acute 2.5 mM mβCD treatment. (**C**) Averaged G-V relationships showing the activation (triangle maker) and inactivation (diamond shape) profiles of endogenous sodium channel before and after acute 2.5 mM mβCD treatment. (**D-E**) Summary data showing the $V_{1/2}$ and the slope factor ($V_s$) for sodium channel activation. (**F-G**) Summary data showing the $V_{1/2}$ and the slope factor ($V_s$) for sodium channel inactivation. Data shown are mean ± s.e.m., n = 5 patches, $*p < 0.05$.

**Supplementary Table 1:** Effects on HCN gating parameters by cholesterol supplementation.

| HCN Gating Parameters | Naïve DRGs | | SNI DRGs | |
|---|---|---|---|---|
| | OMDs | Free Cholesterol | OMDs | Free Cholesterol |
| $V_{1/2}$ | Insensitive | Sensitive | Insensitive | Sensitive |
| $V_s$ (Slope Factor) | Insensitive | Insensitive | Sensitive | Insensitive |
| $\tau_1$ of Activation | Insensitive | Sensitive | Sensitive | Sensitive |
| $I_{max}$ | Insensitive | Sensitive | Sensitive | Sensitive |

*Sensitive* and *Insensitive* indicate whether a given gating parameter is affected or unaffected, respectively, by changes in the ordered membrane domain (OMD) size or free cholesterol following cholesterol supplementation.
